# The Elephant in the room: What can we learn from California regarding the use of sport hunting of pumas (*Puma concolor*) as a management tool?

**DOI:** 10.1101/813311

**Authors:** John W. Laundré, Christopher Papouchis

**Author notes:** Corresponding Author: John W. Laundré, 8070 NE Barberry Dr., Adair Village, OR 97330.

## Abstract

Pumas (*Puma concolor*) in 10 western states of the U.S. have been managed through the use of a sport hunt. The rational for this management technique is that puma populations needed to be hunted to reduce threats to human safety, their livestock, and wild ungulate populations. We evaluated these claims with the state of California as a control, which has prohibited sport hunting since 1972. We tested four hypotheses: 1) Sport hunting reduces puma density, 2) Sport hunting reduces problematic puma-human encounters, 3) Sport hunting reduces puma predation on livestock, and 4) Sport hunting reduces the impact of puma predation on wild ungulate numbers. Results indicated: 1) Puma densities did not differ between California and sport hunting states, 2) California was the 3^rd^ lowest in per capita puma-human incidents. 3) The per capita loss of sheep was significantly lower (*t* = 5.7, P < 0.001) and the per capita loss of cattle in California did not differ significantly, from the other 10 states (P = 0.13). 4). Changes in annual deer populations in California correlated with changes in other states (F = 95.4, P < 0.001, R^2^ = 0.68) and average deer densities in California did not differ significantly from the other states. We concluded that sport hunting of pumas as a management tool has not produced the outcomes sought by wildlife managers and may even exacerbate conflicts between pumas and humans. It is suggested that state agencies re-assess the use of sport hunting as a management tool for pumas.

## Introduction

Pumas (*Puma concolor*), like the other predators in North America, were viewed by European colonialists and their descendants as threats to human safety and domestic livestock as well as competition for wild ungulates, mainly deer (*Odocoileus* sp) and elk (*Cervus elepus*).

Consequently, they were eliminated from much of their range in the Eastern and Midwestern United States by the mid to late 1800’s. In the West, unrestricted trapping and hunting of pumas continued until the 1960’s, with bounties being offered for their removal. By the mid 1900’s, the scientific evidence began to demonstrate the ecological value of large predators, including pumas, in ecosystems [1, 2]. Additionally, many scientists and citizens began to question the ethics of uncontrolled killing of pumas and began to advocate for some degree of protection [3]. However, there remained the perception among wildlife managers that some level of control was still necessary to prevent puma populations from growing to socially unacceptable levels where they might threaten human safety, livestock interests, and population objectives for big game, principally deer [4] (https://idfg.idaho.gov/wildlife/predator-management). In response, in the 1970’s ten of the 12 states where pumas still occurred classified them as a game species and established sport hunting seasons [4]. The two exceptions of this management approach are Texas, where pumas are completely unprotected and can be hunted without limit, and California, where pumas are fully protected from sport hunting and managed through relocation or killing of individuals puma that pose a threat to public safety, livestock or threaten the viability of bighorn sheep populations (https://www.wildlife.ca.gov/keep-me-wild/lion, Accessed on February 28, 2018, https://www.wildlife.ca.gov/Conservation/Mammals/Mountain-Lion/Depredation, Accessed on February 28, 2018).

Though providing some degree of protection to pumas, e.g. closed seasons, the intent of a sport season on puma was to continue to control their populations to address the three main concerns of public safety, livestock and ungulate protection [5, 6]. As a result, the primary management objective (MO) for sport hunting of pumas in these ten western states was to usually set “bag limits” similar to historic bounty kill levels, which never exceeded 1,000 animals per year.

However, since the enactment of sport hunting, the number of pumas killed annually by sport hunters has steadily increased. By 2016, the 10-state average kill rate of pumas was 390 per state or over 3,900 individuals per year (Fig. 1). Of these, 3400, or > 89%, are killed by sport hunters and the rest for specific threats to human safety, livestock depredation, or accidents (Unpublished agency reports). This sustained high-rate of puma killing has elicited questions as to whether sport hunting actually achieves its purported management goals [7].

**Fig. 1.**
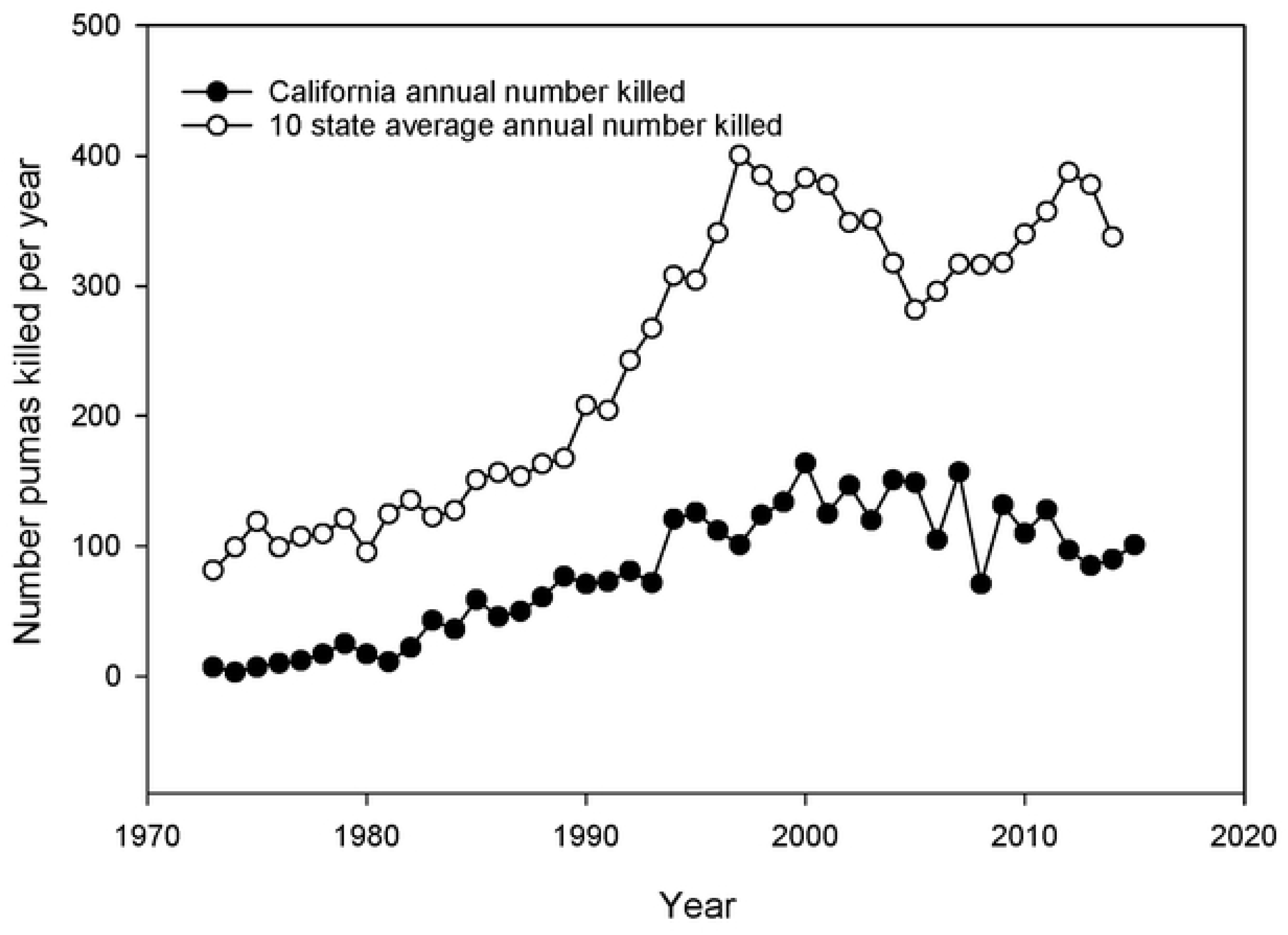
Number of puma killed per year in California compared to the mean for the 10 western states with a sport hunt of puma. The numbers for California represent animals specifically identified as conflicting with human safety or livestock depredation and other causes. The numbers for the 10 other states represent animals killed by sport hunters (80-90%), ones specifically identified as conflicting with human safety or livestock depredation, and other causes.

Most state game agencies rely on the North American Model for Wildlife Conservation (NAM) for guiding their management policies [8]. The NAM explicitly advocates the use of science and research in setting and justifying wildlife management policy [8,9,10,11]. Nevertheless, one recent evaluation of hunt management in the United States and Canada found little adherence to science-based approaches [12]. Using the criteria of Artelle et al. [12], our assessment of available state management plans for pumas, indicates this to be the case for pumas in most of the western states. Additionally, there have been more recent calls for using science to evaluate the possible politicization of wildlife management decisions [13].

Regarding puma management, what does the science tell us? An increasing number of scientific studies have questioned the putative effectiveness of sport hunting to meet MO’s of state agencies. Specifically, sport hunting of pumas might not reduce puma numbers [14], or result in larger ungulate populations [15,16,17,18]. Several studies provide evidence that sport hunting increases the rate of puma interactions with people and livestock, thereby exacerbating the very problems it is intended to ameliorate [5,19, 20]. This growing body of data has placed doubt upon whether sport hunting is an effective management tool for pumas.

Employing the guidelines of adaptive management [4], it is appropriate to ask whether sport hunting has been successful in meeting management objectives over the ∼ 45 years since it was initiated in the western U.S. In doing so, we identified four hypotheses that emerge from desired outcomes articulated by state management agencies. These are that 1) sport hunting will suppress puma populations, 2) sport hunting will reduce the number of problematic puma-human encounters; 3) sport hunting will reduce puma predation on domestic livestock, and 4) sport hunting will reduce the impact of puma predation on wild ungulate numbers, resulting in increased hunting opportunities for the sport hunt of ungulates.

Unfortunately, these hypotheses are difficult to test. In particular, as each of the 10 states have continued to rely on this management strategy, there is no “control” within or among those states other than to alter the number of pumas removed by the sport hunt. One state, Washington initiated a metapopulation style management program [21] where levels of killing of pumas were specifically set for designated management units [22]. As a result, it is in this state that researchers have been able to test some impacts of the sport hunt with the previously mentioned contradictory findings [5,14, 20]. However, except for these localized within state comparisons, we are not aware of any large scale, multi-state test of the sport hunting hypotheses.

Fortunately, the state of California offers a potential control for such a multi-state test. California has not used sport hunting to manage pumas over the same time period the other states have employed it. Instead, since 1972, California has handled puma-human conflicts and livestock depredation on a case-by-case basis and specifically removes animals causing these conflicts. There is no killing of pumas specifically with regards to management of wild ungulate populations, except for threatened bighorn sheep (*Ovis canadensis*). As a consequence, over the same 45 years, the number of pumas killed in California has been consistently lower (< 150 animals/year) than those states with sport hunting seasons on pumas. Thus, California would appear to be an appropriate “control” to compare against the “treatment” of a sport hunt. Since the remaining 10 states have sustained some level of sport hunting as a management strategy used over the time period, this comparison should enable a test of whether a sport hunt management strategy is achieving desired management goals.

The predictions specifically are that California, in the absence of a sport hunt of pumas should have 1) higher puma population densities; 2) a higher percent of problematic puma-human encounters; 3) higher percent of puma predation on domestic livestock; and 4) higher levels of puma predation on ungulate populations, resulting in lower hunting opportunities for sport hunting of ungulates, specifically deer. If these predictions are supported by the 40+ year data base available, then it would lend support to the hypothesis that sport hunting of pumas is a reasonable management strategy to obtain the desired results as stated above. If these predictions are not supported, then it would be reasonable to reject this hypothesis.

## Methods

### Study Areas

The 10 western states that use the sport hunt management strategy encompass most of the diverse habitat types found in the Western United States. Pumas are found throughout most of these habitats but are rare in some of the harsher, dryer areas of each state. As a result, puma range in most states is less than the total area of the state. Most states have estimated the suitability and extent of different puma habitats in their states. Where state estimates were not available, we used recent data based on GIS analyses conducted by the Humane Society of the United States [7] (HSUS). As on average, HSUS habitat estimates only differed from state ones by approximately 4%, HSUS estimates were considered reliable enough to use when state estimates were lacking. Each state agency also has estimates of the amounts of appropriate deer (mule *O. hemionus* and white-tailed *O. virginianus*) habitat occurs within their boundaries. In most cases, puma and deer distributions overlap. California, which extends from the border with Mexico north to Oregon, contains most of the major ecosystems found in the West, from desert to high mountain forests [23]. As such, the impact of habitat differences on comparisons between California and the other 10 states could be considered minimal. The state also has identified the amount of appropriate puma and deer habitat. We used the estimates of total area of habitat for pumas and deer from each state when making density calculations.

### Data sources

For all the comparisons made, we relied on data sets generated by either state or federal agencies or in the case of puma-human incidents, private organizations/individuals. These data sets have been maintained and published as open public records. We recognize that the reliability and scientific rigor of these data has been questioned. However, we argue that any testing of the sport hunting hypothesis should be done with the same data used to justify sport hunting as a management tool. We further argue that if these data are not considered rigorous enough to test these hypotheses, then they should not be used in making management decisions. However, many of these data sets, e.g. deer/puma population estimates and livestock depredation estimates, are routinely used by state agencies in their management decisions, consequently, we used them to test the hypotheses regarding sport hunting presented here.

State and Federal data sets used in our analysis include 1) estimates of puma abundance, 2) numbers of pumas killed yearly by sport hunters and other causes, 3) estimates of deer populations, 4) estimates of the number of deer killed yearly by hunters, 5) estimates of the inventory of livestock, cattle and sheep, and 6) estimates of the number of livestock, cattle and sheep killed by pumas. Estimates of the number of puma-human incidents for each state have been maintained mainly by individuals and published either in the scientific literature [24] or available on the internet (http://tchester.org/sgm/lists/lion_attacks.html). These estimates were cross checked with inquires to state agencies as to records they had and updated as necessary.

In making comparisons, we first designated three basic stages in the evolution of the sport hunt of pumas. These are our designations based not on recognized agency policy but on our interpretation of documented puma population and sport kill data available. The first 20 years (∼1970 – 1990) we refer to as the recovery period as puma populations were presumably still low from the decades of uncontrolled killing and the reported killing of animals by sport hunters was also low (∼ 100-150 per state per year; Fig 1). By 1990, various studies indicated that puma populations in general had recuperated (the recovered period, 1990-1999) and were increasing and decreasing with available resources [25, 26]. The sport killing of puma was beginning to increase during this time and along with other human sources of mortality peaked at around 400 per year per state in 2000, with 88% being from the sport hunt (Fig. 1). From approximately 2000 to 2015 (the intense management period) total mortality of pumas remained between 300-400 animals per state per year, again 80-90% from the sport hunt. As puma populations and kill rates were low during the recovery period for the 10 states, inclusion of this timeframe in comparisons might dilute effects of the sport hunt on the metrics we compared. Thus, most of our comparisons covered the last two periods as any effect of sport hunting should be more prominent, especially during the last 15 years of intense management.

### Standardizing the data

Because the data used for deer and pumas come from a wide geographical area and at least 11 different governmental agencies, we attempted to standardize the data in several ways. Most estimates of abundance or kill levels of deer and pumas were converted to population densities or kill densities (number killed/habitat area) based on the aforementioned estimated areas of appropriate habitat. Kill (= harvest) densities are commonly used by state agencies to set MO’s for puma kill limits. Kill densities for puma were per 10,000 km^2^ while kill densities for deer were per 100 km^2^. In some instances, we converted individual entries of a data set to the percent they were of the maximum entry of that data set. This “percentage of the maximum” facilitated comparing patterns of change as well as amplitude of that change among the diverse data sets.

Estimates of puma mortality by all sources come from records maintained by state agencies. Total mortality levels were primarily (> 80 %) from sport hunting in the 10 states under consideration. However, as the level of mortality from California was just from all other causes, in making our comparisons we used the total number of pumas killed in a state rather than just the number killed by sport hunting. Also, some states include non-hunting deaths of pumas in setting their MO’s.

To standardize livestock data across states, we converted the estimated number of animals killed by pumas to the percentages they were of total head inventory exposed to predation, e.g. livestock on open range. These data were retrieved from appropriate USDA documents (https://www.nass.usda.gov/, Accessed on February 28, 2018). In these documents, total cattle inventory of a state included beef and dairy cattle. We subtracted the number of dairy cattle from the total to obtain an estimate of the number of beef cattle, animals most likely to be grazed on open range. There was a category of cattle on feed (= feedlots), but because these cattle could have come from anywhere, including other states, we did not use these estimates to adjust the inventory of beef cattle in a state. Consequently, we assumed all beef cattle were at least at sometimes grazed on open pasture exposed to possible predation by pumas. Data on calves were separately available. We did not use inventory data on cattle in Texas because most of the beef cattle in Texas are raised outside of current puma range and there were no estimates available for the number of beef cattle in the proportion of Texas where pumas occurred [27].

National levels of cattle and calf losses to predators, including pumas were reported yearly. However, there were only 5 years (1991, 1995, 2000, 2005, and 2010) where those losses were separated out by state and cause specific by predator (http://usda.mannlib.cornell.edu/MannUsda/viewDocumentInfo.do?documentID=1625, Accessed on February 28, 2018). Thus, we used only these 5 years in our comparisons of per capita cattle and calf loss by pumas in California verses the other 10 states.

For sheep and lambs, the data categorized all sheep as sheep and lambs combined and also reported the annual lamb crop. The lamb crop was not identified as before or after docking but we assumed it was the same for all states. Because simply subtracting the lamb crop from the total sheep did not always provide us with credible estimates for adult sheep only, we used the categories of “all sheep” (adults and lambs) and “lamb crop” in our comparisons. We assumed all sheep and lambs were at sometimes grazed on open range and thus exposed to possible predation by pumas. We included Texas in some of the comparisons of levels of puma predation on sheep and lambs because most sheep in Texas are raised within current puma range in that state [27].

Annual losses of sheep and lambs to predators, included pumas, were available yearly but there were only 5 years (1990, 1994, 1999, 2004, and 2014; http://usda.mannlib.cornell.edu/MannUsda/viewDocumentInfo.do?documentID=1628, Accessed on February 28, 2018), where the data were separated out by state and cause specific by predator. Thus, we used only these 5 years in our comparisons of percentage of sheep and lamb loss by pumas in California compared to the other 10 states.

Attack and mortality data from puma attacks on humans were available from before 1900. However, as we were interested in the risk of humans since the early 1970’s, we only used the data compiled since 1972, specifically during the recovered and intense killing periods. We estimated per capita (per million people) attack and mortality rates based on total human population estimates within each state for 2010, year of last census. As pumas are widely distributed over most western states and known to use exurban and suburban areas and many urban persons visit areas where pumas are, we used the total populations reported for each state. Texas, however, was excluded from any analysis of per capita incidents because we could not find population data on just the region of Texas where pumas occur. To include the more populated portion of the state where pumas were not found, would bias any comparisons.

When comparing deer data among states, we standardized the data relative to density (#/km^2^). As deer abundance is estimated in similar ways across states, e.g. aerial surveys, we assumed the values reported, converted to densities, could be comparable across states. There could be some inherent differences in possible densities based on the proportion of habitat quality within a state, e.g. desert shrubland versus high altitude alpine vegetation. We address the effects of these differences in the discussion of the results of our comparisons. Deer population densities and deer kill densities (by hunters) were calculated with agency published estimates of deer habitat within each state. Hunter success and the number of deer per hunter were calculated based on the number of licenses sold. In some cases, we again further standardized the data as percentages of the maximum value recorded to facilitate comparisons of trends.

In all the comparisons, data from California were evaluated directly to the equivalent data of the 10 states with the sport hunt of pumas. Under this design, when appropriate, a t-test or its non-parametric equivalent for a single observation compared to a sample was used. In the case of any correlation analyses, any comparisons of correlation coefficients were made with appropriate statistical tests. If percentages were compared, they were first transformed with the recommended arcsine square root transformation [28]. Again, we recognize that others have argued that some of the data collected by agencies may not withstand the rigor for statistical analyses. However, we again argue that these are the only data available and are used by agency scientists in their analyses and decision making. As it are these data that the entire hypothesis for sport hunting rests, it should be these data that are used for the testing of that hypothesis.

## Results

### Prediction 1: California will have higher puma population densities

We first tested whether sport hunting has led to reduced puma populations or at least keep them lower than in the absence of a sport hunt (California). Puma are notoriously difficult to enumerate. However, all game agencies have at one time or another published estimates of puma numbers within their state. These estimates can vary widely and high and low values are usually given. Unfortunately, the years of these estimates across states rarely coincide. For 2003, however, most agencies provided high and low estimates for pumas in their state [29]. As these estimates were provided after over 30 years of control (California) and treatment (10 states with sport hunt management), it would seem reasonable to compare densities between these states and California. We selected the high estimates as these values are commonly the default numbers cited by agencies when developing of management guidelines (Fig 2). As can be seen in Fig. 2, estimates of puma densities in California are not higher than, but are rather at the average of those states with sport hunting. Thus, the data do not support the hypothesis that after 30+ years, puma densities in states with sport hunting of pumas are significantly lower than in California. In fact, half of the sport hunting states reported puma densities higher than California.

**Fig. 2.**
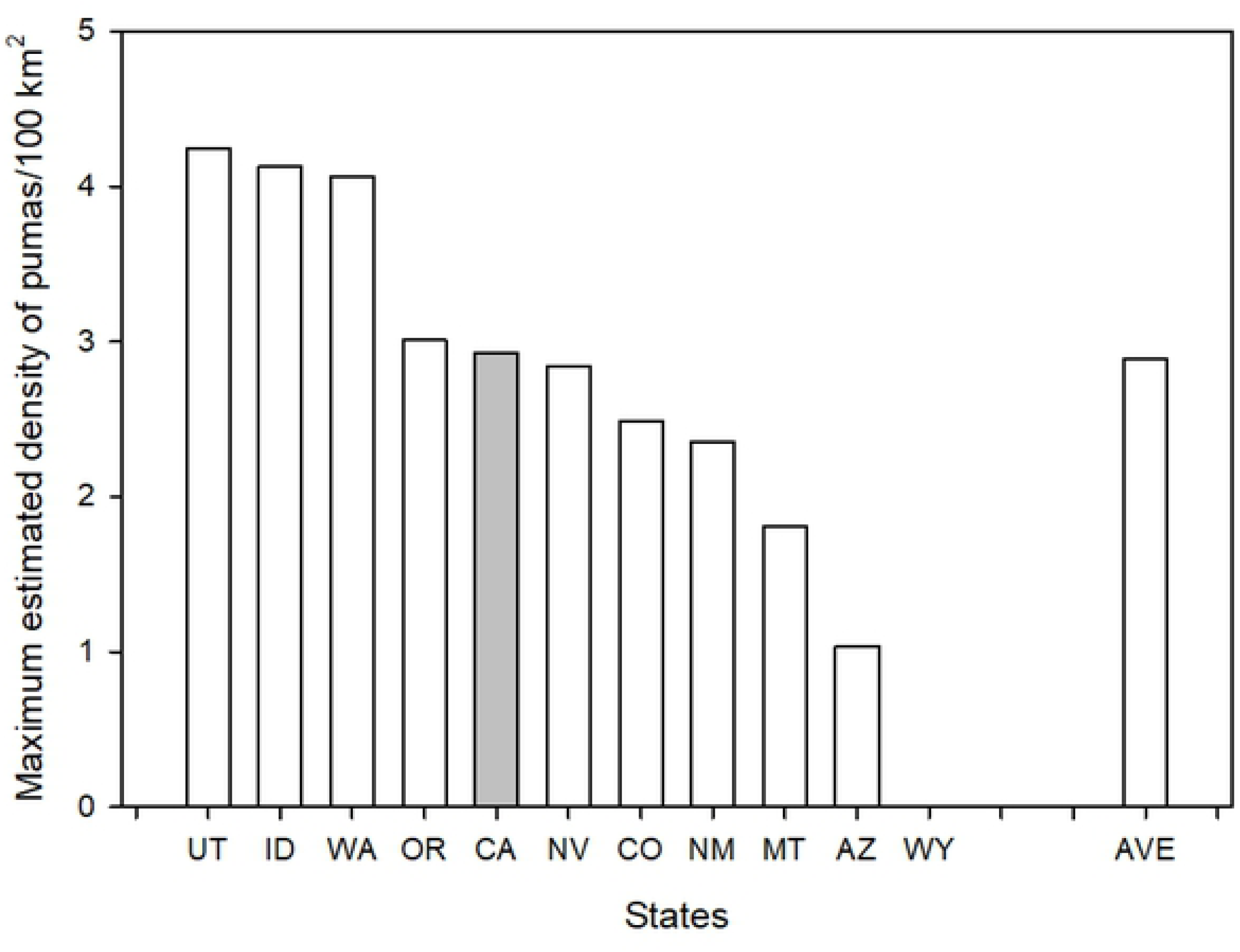
Maximum estimated density of puma (animals/100 km^2^) in 2003 for 9 of the western states with a sport hunt of puma (no estimate was available for Wyoming) and California (dark column). Estimates are based on data provided by agencies in Becker et al. (2003) and agency reported amounts of puma habitat within their boundaries. States are identified by their standard two-letter postal codes. The final column (AVE) is the average for the 9 states that have a sport hunt of puma.

Additional comparisons can be made for any of the states where later estimates are provided. The prediction is that after 12 years of intensive sport hunting, estimated puma population densities within a state should be lower than the 2003 estimate, while California should have no difference. California currently lists its mountain lion population to be between 4,000 and 6,000 animals (https://www.wildlife.ca.gov/Conservation/Mammals/Mountain-Lion/FAQ#359951241-how-many-mountain-lions-are-in-california, Accessed on February 28, 2018), which is the same reported for 2003. Arizona currently states it has between 2,500 and 3,000 pumas, placing the current maximum number 500 above the maximum reported in 2003. Montana reports a 2017 maximum estimate of 5,000 pumas [30], which represents a state-wide density of 2.8 animals/100km^2^. Though there is no earlier statewide estimate to compare against, this density is only slightly below the 3.27 animals/100 km2) reported in one study area in western Montana [31], suggesting little change in total numbers since that time. New Mexico reported estimated its puma population at 3,123-4,269 animals in 2017 [32], a > 45% increase from the 2003 estimate of 2,150 animals. In sum, population estimates provided by agencies do not depict declining puma numbers in states with the sport killing of pumas. Over the same period, puma numbers have not reportedly increased in California where they are protected from sport hunting.

Oregon, bordering California to the North, has published estimates of puma numbers since 1994. These estimates have been used by the Oregon Department of Fish and Wildlife to guide its puma management decisions, including sport hunting mortalities, which have steadily increased since 1994 (Fig. 3). We compared ODFW’s puma annual population estimates with puma mortality levels and found a significant positive correlation (P < 0.001, R^2^ = 0.74). In effect, it appears that the more animals that are killed in Oregon, the higher the reported population. This is the exact opposite that is predicted by the sport hunting hypothesis.

**Fig. 3.**
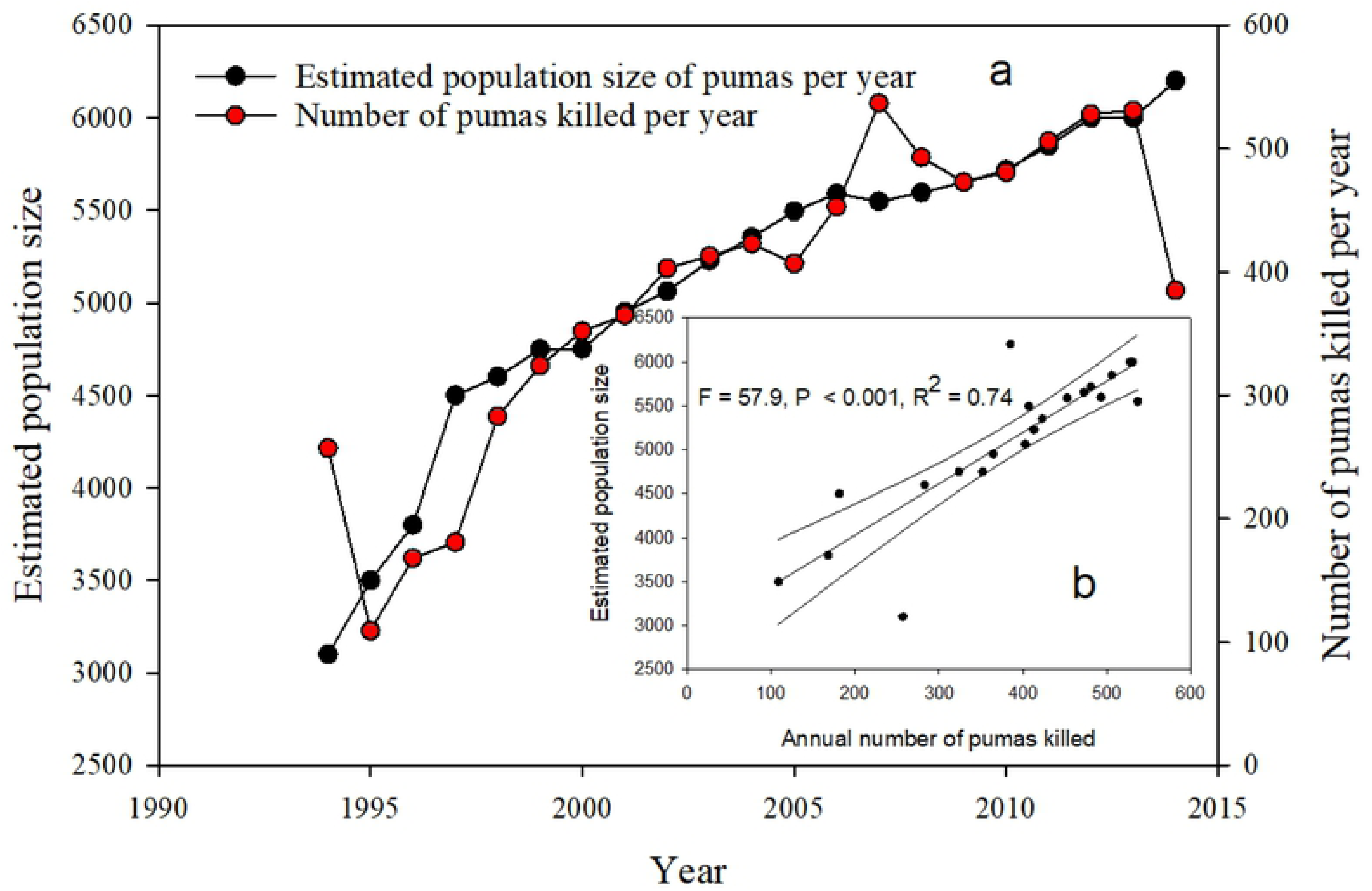
(a) Estimated population size and number of puma killed per year in Oregon as reported by Oregon Department of Fish and Wildlife (http://www.dfw.state.or.us/wildlife/cougar/) for 1994 to 2014. Fig. 3b is the correlation of estimated population size with annual number of pumas killed.

In sum, based on the available data, we found no support of the hypothesis that sport hunting controls puma numbers below the level expected in the absence of this management practice.

### Prediction 2: California will have higher number of per capita puma-human incidents

The test of the sport hunting model for this prediction is whether or not states using this management technique are experiencing fewer problematic puma-human interactions than California. We compared the per capita (per million persons) number of puma attacks and human fatalities that have occurred in California to the 10 states with sport hunting. The few overall numbers of such mortalities over the last 100 years makes statistical comparisons difficult.

Consequently, we used the combined non-fatal and fatal attack data in our comparisons. In sum, as of 2016, 84 puma attacks on humans, non-fatal and fatal, have been recorded since 1972 (beginning of sport hunting) in the twelve western states (Fig 4a). Most states reported 5 or fewer incidents over the 44 years. The highest was Washington with 16, followed by California with 15 and Colorado with 13. Texas and Arizona each reported 8 incidents. On a per capita basis (per million persons), California ranked 3^rd^ lowest with 0.40 attacks/million persons whereas Montana was highest with 7.1/million persons. The pattern does not change, including in reference to California, when we considered the time span of 2000-2015, the period of increased killing of pumas by sport hunting (Fig. 4a).

**Fig. 4.**
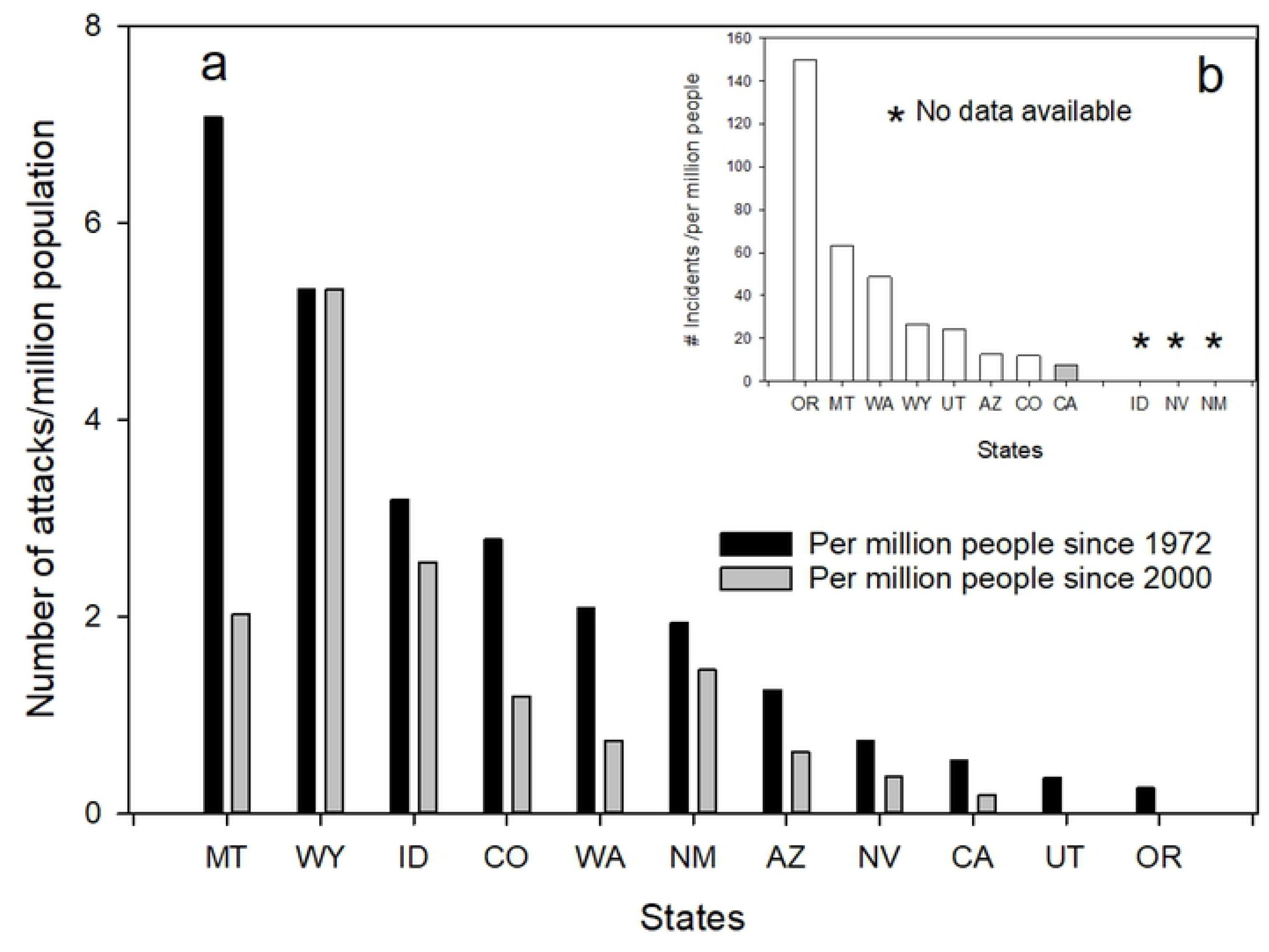
(a) per capita (per million humans) of cougar attacks on humans for the 10 western states with a sport hunt of puma and California. Per capita rates are based on total population (2010 census) of states. Fig. 4b is per capita rate of cougar-human incidents, including attacks, threats, and livestock depredation for the 8 states reporting these data. Idaho, Nevada, and New Mexico do not maintain records of incident reports. States are identified by their standard two letter postal code.

Another indicator of puma-human conflicts is the number of incidents reported per year. California and seven of the 10 states with a sport hunt, recorded incidents that they considered as being serious enough to respond to (Fig. 4b). Some of these were actual attacks but many involved perceived threats to person or pets or livestock. California reported an average of 200 incidents/yr since 2000. Though most of the states that use sport hunting had fewer than 100 incidents, Washington (578/yr) and Oregon (328/yr), reported higher numbers of incidents than California. However, again, on a per capita basis, California ranked the lowest of the states reporting (Fig. 4b).

Annual incident data were available from the early-mid 1990’s to 2018 for California and three other states (Oregon, Utah, and Washington). When we correlated puma kill density rates with incidents for these 4 states, there was no correlation for California and Oregon but there were positive correlations for Utah and Washington, indicating higher puma kill rates coincided with higher number of incidents (S1 Fig.).

Based on the attack and incident data, we found little support for the hypothesis that the sport hunting of pumas decreases the level of risk humans faced from pumas.

### Prediction 3: California will have higher percentage of puma predation on domestic livestock

Besides human safety, the second most frequently offered rationale for sport hunting of pumas is that it should reduce incidents of livestock depredation, principally cattle and sheep. To test this prediction, we used cause-specific depredation rates by pumas on livestock and compared among states the percentage loss from pumas based on the total number of head exposed to possible puma predation (see Methods for details). We present means for the specific years when cause specific predation was reported.

### Cattle

Overall cattle losses to pumas are extremely low, less than 0.2% of total head inventory. Figure 5a ranks the 11 states relative to the average percentage of cattle lost to pumas during the 5 years reported (see Methods). California reported higher percent cattle losses than 8 states and lower losses than two states (Fig. 5a). These patterns were similar for calves (Fig. 5b). In comparing the percentage loss for the 5 years examined (See Methods) between California and the average loss for the other 10 states, there no significant differences for either cattle (paired – *t*, P = 0.56) or for calves (P = 0.132).

**Fig. 5.**
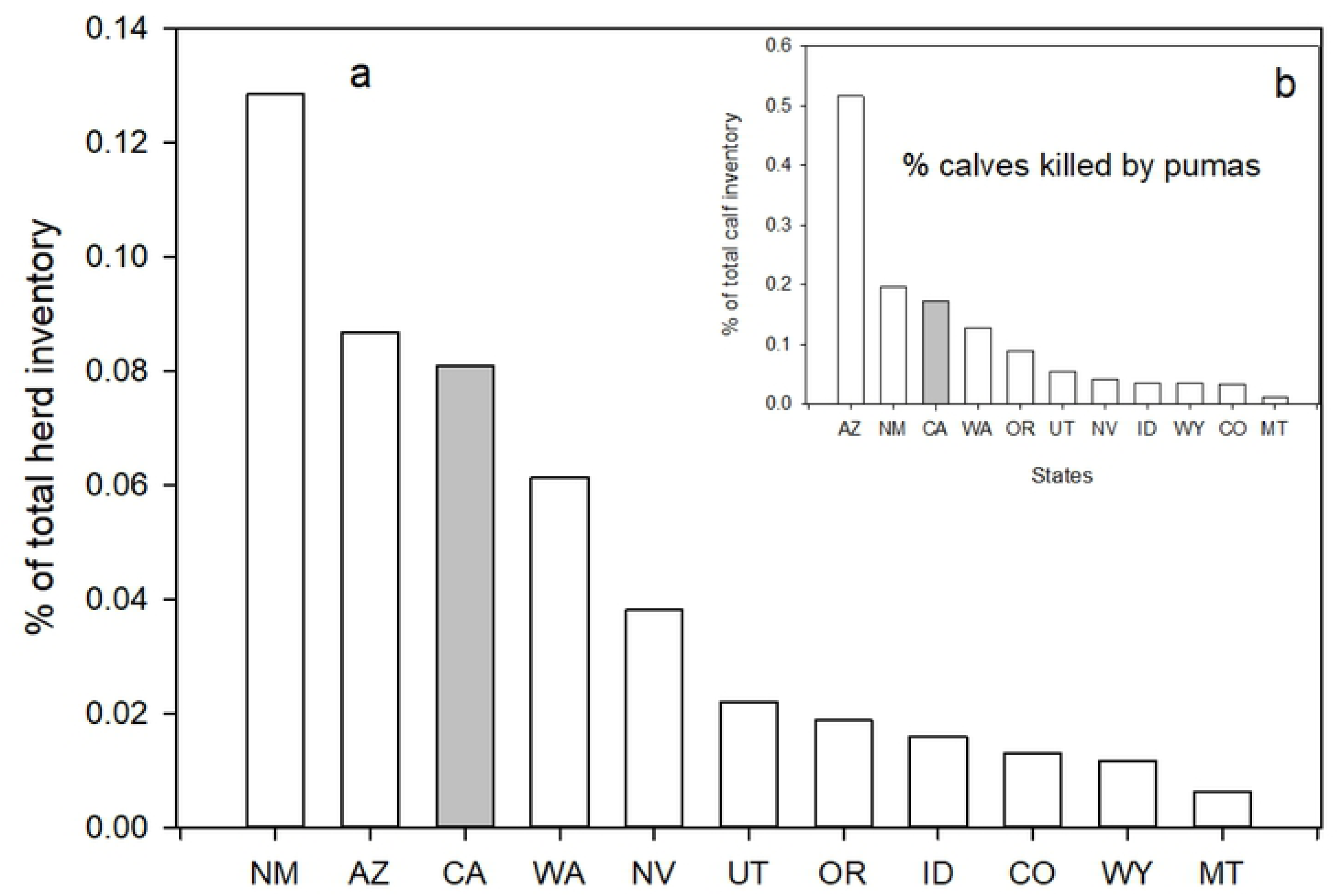
Per capita (percent of total available herd inventory) predation of puma on cattle (a) and calves (b) in the 10 western states with a sport hunt of puma and California. States are identified by their standard two letter postal code.

To further test whether sport hunting reduced cattle losses, we combined the data for percentage loss of calves from the 10 states with a sport hunt for the 5 years where data were available and correlated them with the puma kill density for the years previous to the sample years (Fig. 6a). Kill density of pumas was used to standardize the mortality rate across states. The prediction tested was that the percentage loss of calves would be negatively correlated with the number of pumas killed the previous year. The correlation was not significant (Fig. 6a). When we added the California data to the graph, but not the correlation, California had the lowest per area kill rates and also some of the lowest percentage loss of calves (Fig. 6a). The same analysis using cattle lost also showed no correlation.

**Fig. 6.**
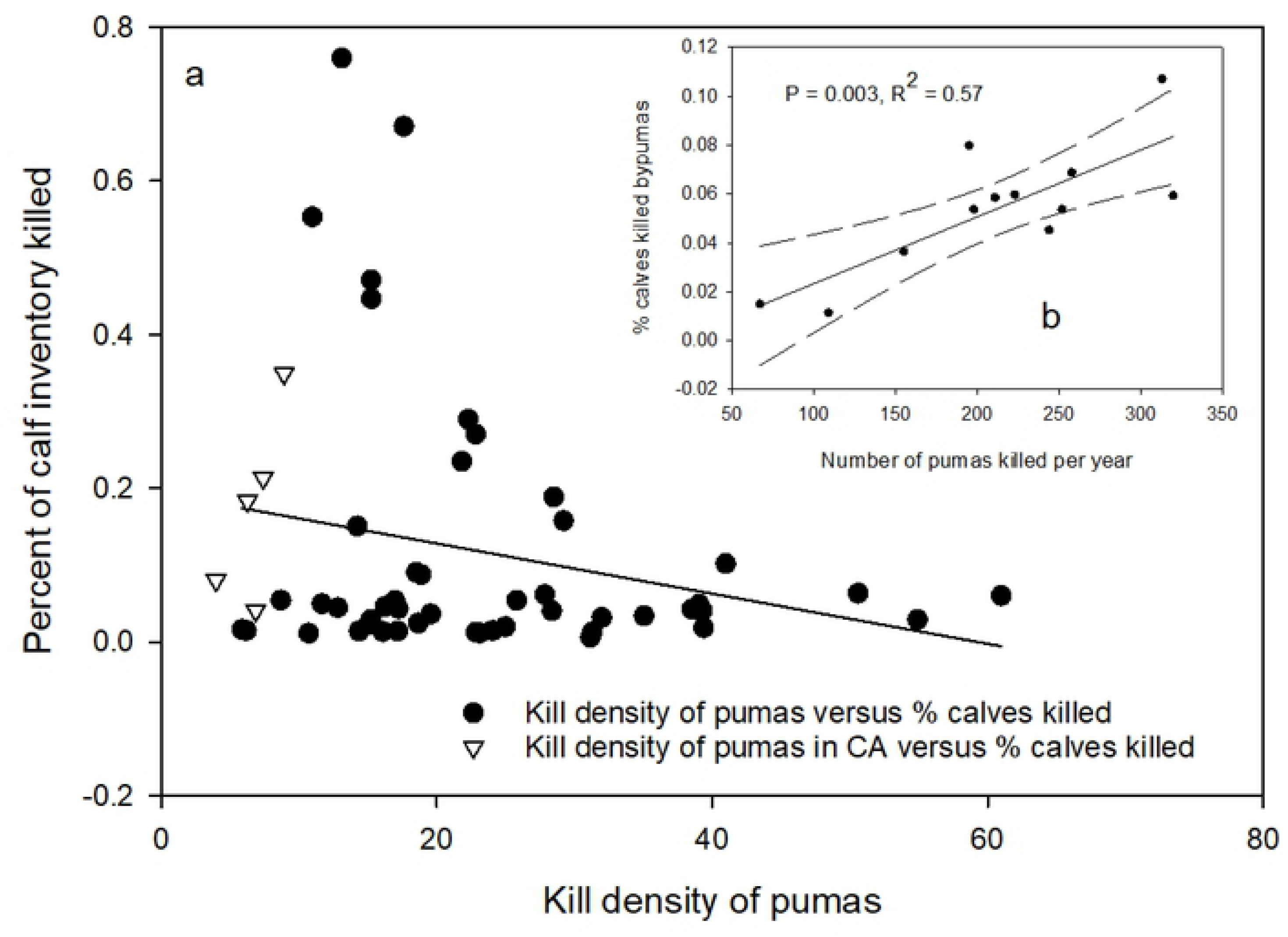
(a) Correlation of percent calves killed by puma with puma kill density (# of puma killed per 10,000 km^2^ of habitat) for combined data from 10 states with a sport hunt on puma. Data from California are included in the graph for comparison but were not included in the correlation analysis. Fig. 6b is correlation of percent calves killed by pumas in Wyoming with number of pumas killed per year for 2004 to 2012. Data are from the 5 years where cause specific predator mortality were available (1991, 1995, 2000, 2005, & 2010).

One state, Wyoming, maintained cause specific depredation records for multiple years, including annually from 2004 to 2012. For each of those years, we compared the number of pumas removed the previous year with the percentage of cattle and calves killed for each year. The prediction is that if sport hunting puma is beneficial to cattle survival, there should be a negative correlation between the number of pumas removed one year and the percentage loss of cattle and calves the following year. The results indicated no relationship between cattle loss and puma kill rates. However, calf losses were positively correlated with the number of pumas killed the preceding year (Fig. 6b, P = 0.003, R^2^ = 0.58). Higher calf losses were associated with higher numbers of puma killed, contrary to the prediction.

### Sheep

The livestock inventory data did not clearly differentiate sheep and lambs but did present estimates for lamb crops. Sheep losses by pumas however, were clearly indicated as either adult sheep or lambs. As the inventory data were often incompatible, e.g. total sheep minus lamb crop did not equal an estimate of adult sheep, we only compared total puma depredation losses as a proportion of combined sheep and lambs and then puma depredation on lambs as a proportion of the lamb crop. For all states, including now Texas, data were available only for specific years (See Methods) and so we present the means over those years.

Relative to the mean percentage of inventory of all sheep and specifically for lamb losses over the 5 years where data were available (See Methods), California ranked 6^th^ of the 12 states (Fig. 7a & b). When considering the 6 years separately, the percentage lamb loss to pumas in California for each year was significantly lower than the corresponding mean for the other 11 states (Paired *t* = 3.53, P = 0.0077). This was also the case for all sheep combined (Paired *t* = 5.692, P < 0.001).

**Fig. 7.**
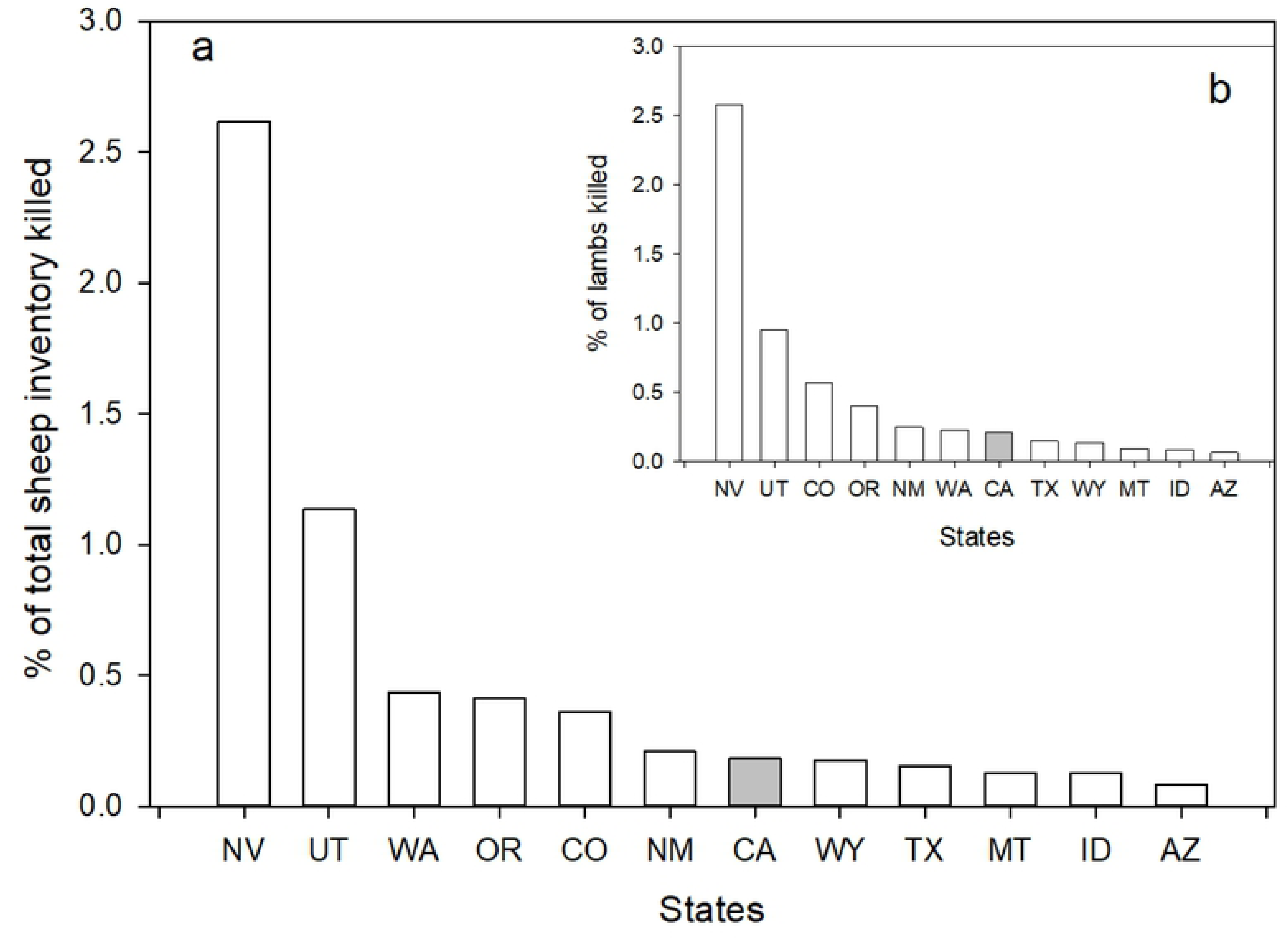
Ranking of each western state with puma (Texas included) relative to percent of total sheep (a) and total lambs (b) killed by pumas. Percentage of animals lost per state were means of the 5 years data were available (1990, 1994, 1999, 2004, & 2014).

As with cattle, we combined the total sheep loss data from the 10 states with a sport hunt for the 5 years and correlated them with puma kill density for the years previous to the sample years. Again, there was no significant correlation. When we added the California data to the graph, but not the correlation, California again had some of the lowest percentage loss of sheep per number of puma killed. When we repeated this analysis for just lambs lost, again, no correlation was found.

Three states, Wyoming, Colorado and Utah, maintained cause specific losses of sheep and lambs to pumas for multiple years and we correlated sheep and lamb losses for those years with the level of puma killed for the years before. None of the correlations for Wyoming and Colorado were significant. For Utah there was a significant (P = 0.05) positive relationship between the number of pumas killed the year before and the percentage of lambs lost and the correlation explained 16% of the variation seen. When all sheep losses were correlated with puma mortality levels, again the relationship was positive, significant (P = 0.049) and explained 16% of the variation seen.

The results of the comparisons of livestock losses from pumas did not support the hypothesis that sport killing of pumas resulted in lower per-capita losses of cattle or sheep. In point of fact, in a few cases, the exact opposite of what was predicted was found: higher mortality rates of pumas were correlated with higher losses of livestock.

### Prediction 4: California will have higher puma predation on ungulate populations, specifically deer

Several metrics are available to test the prediction that killing of pumas via the sport hunt will enhance deer populations or hunting opportunities for hunters. Two useful metrics are estimated deer density and deer hunter kills. These records are maintained by state agencies and commonly used as indicators of population trends (http://cpw.state.co.us/thingstodo/Pages/Statistics-Deer.aspx, Accessed on February 28, 2018). We used both metrics in the following comparisons. As explained in the Methods, we primarily limited our analyses to two timeframes: 1991-2015 and from 2000-2015.

We initially compared long term pattern of changes in deer kill densities from 1927 to 1972 between California and the average for 3 states that also had these data sets (Arizona, Oregon, and Utah). We sought to determine if California had any inherent differences in changes in deer abundance before the sport hunt of pumas was initiated relative to other states, which might affect any comparisons over later timeframes. We standardized the data by calculating the percentage each year’s estimate was of the year with the maximum estimate recorded, which would equal 100%. This allowed us to more directly compare patterns of change in deer kill densities.

When we compared the percent of the maximum deer killed for each year for California and the three-state average for the other states, we found a relatively high degree of concordance (Fig. 8b). Based on kill records, all deer populations experienced exponential style growth in the 40’s and 50’s, peaking around 1960. After 1960, deer populations of California and the other three states appeared to decline in a similar pattern. When compared with a simple correlation of the transformed percentages, the correlation was highly significant (Fig. 8b; F = 95.4, P < 0.001, R^2^ = 0.68). Thus, as indicated by annual kill levels by hunters, changes in California’s deer population before the beginning of the sport hunt of pumas appear comparable to other western states. At times, the magnitude of changes was different but the pattern of change matched. Consequently, any difference between California and the other states during the period of the sport hunt in those states could then be more likely because of the management differences.

**Fig. 8.**
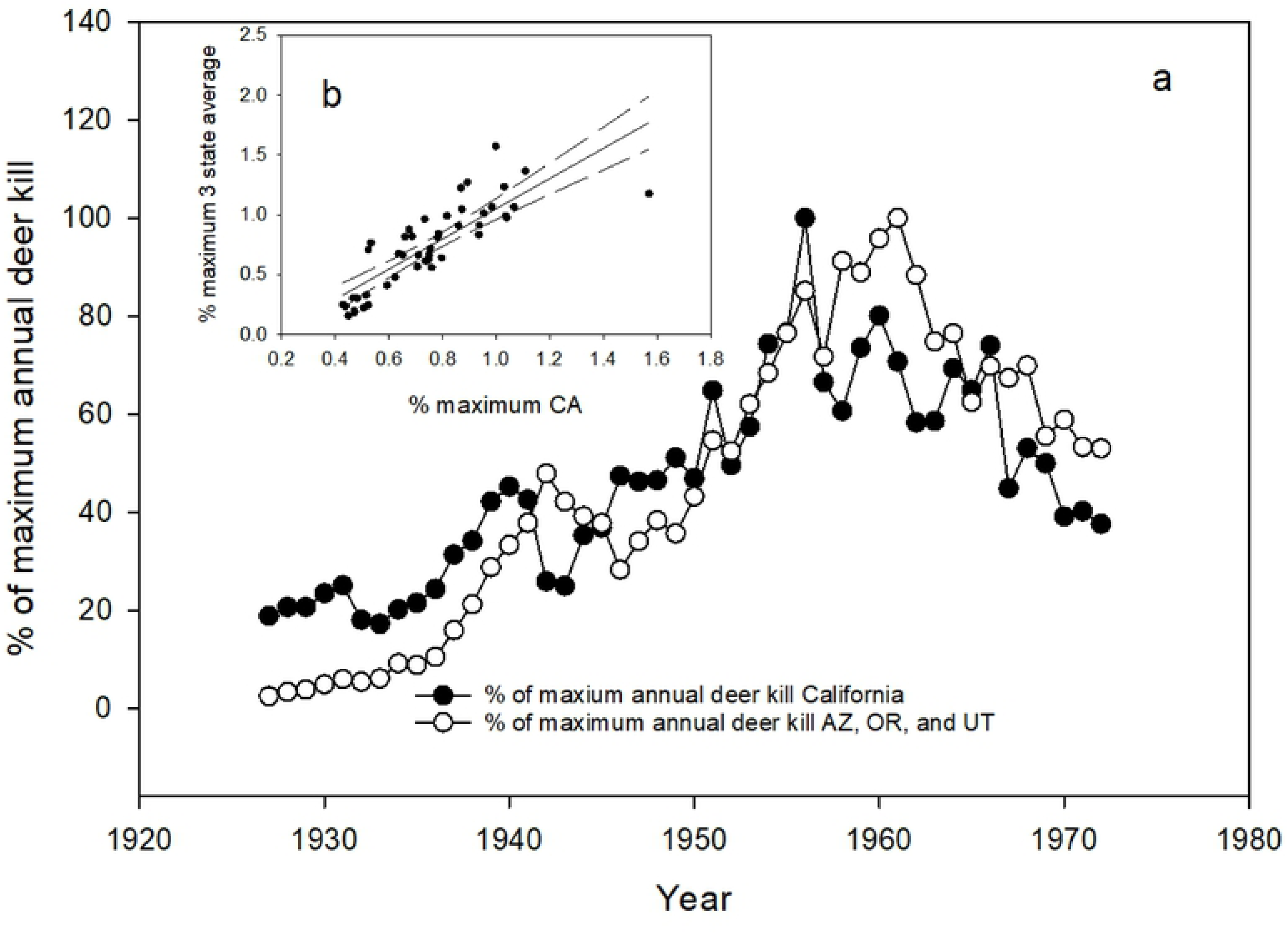
(a) Number of deer killed by hunters each year (1927-1972) expressed as a percentage of the year with the highest deer kill level for California and the average for Arizona, Oregon, and Utah. Figure 8b is the correlation of the mean percent of maximums for the three states versus percent maximum for California.

To test for those differences in these later time periods, we compared California to the 10-state average from 2000-2015 (Fig. 9a). Many states did not have kill data back to 1990 and so we limited our comparison just to the later timeframe of most intense puma kill rates. The prediction tested was that California should exhibit different patterns of change than the other states. For these comparisons, we also converted the number of deer killed in each year to percentages of the year of maximum annual deer kill within that timeframe, to make the lines more comparable. We found (Fig. 9a) again that the patterns of change in deer kill density for California matched closely the pattern of the average for the 10 states. Of note is that California and most of the other states experienced increased deer kills within the last 4 years, supporting the reported estimates of increasing populations of deer in most western states [33, 34]. We found that these data were also significantly correlated (F = 19.1, P < 0.001, R^2^ = 0.55, Fig. 9b).

**Fig. 9.**
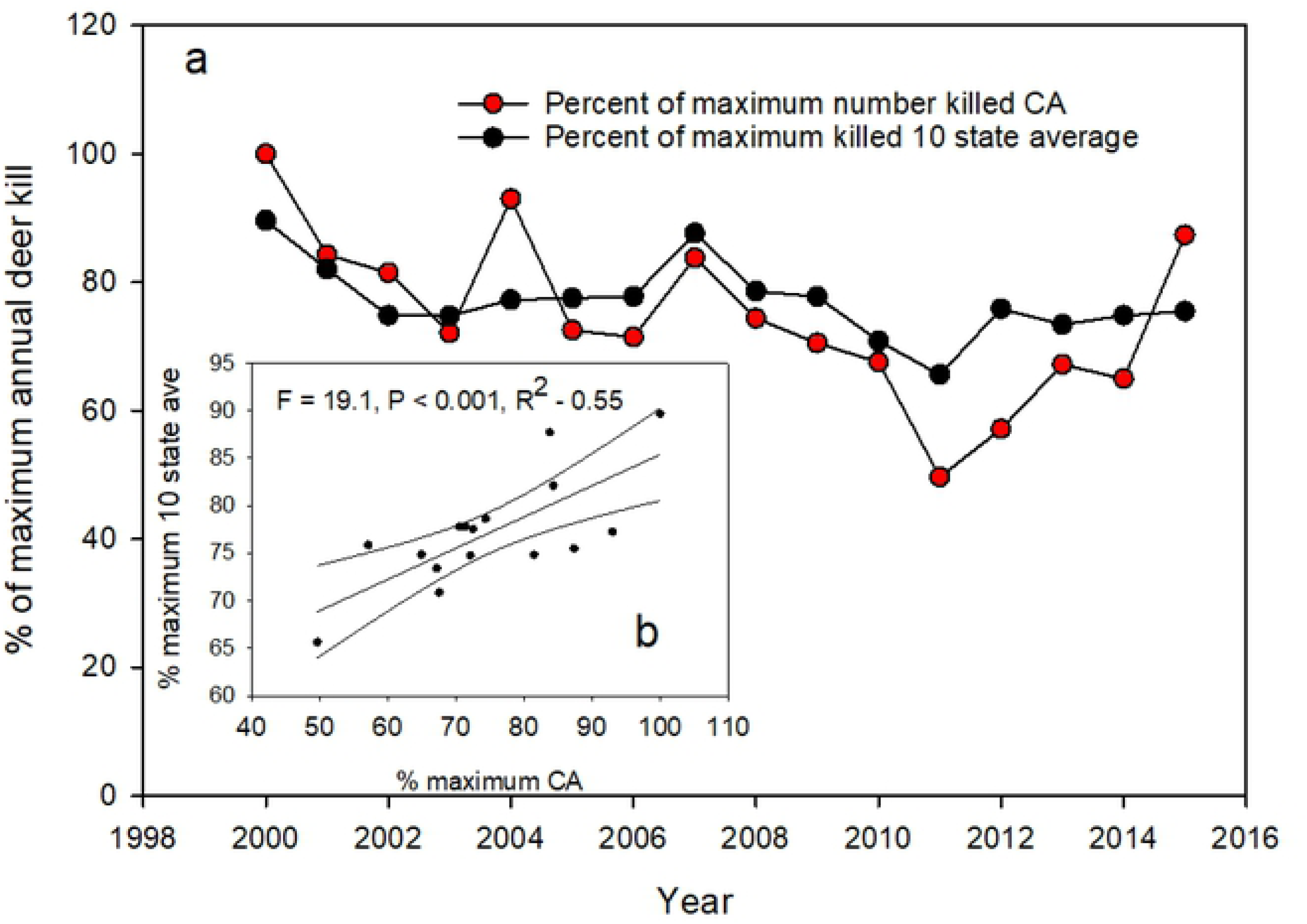
(a) Number of deer killed by hunters each year (2000-2015) expressed as a percentage of the year with the highest deer kill level for California and the mean for the ten states with a sport hunt of pumas. Figure 9b is the correlation of the mean percent of maximums for the 10 states versus percent maximum for California.

As deer populations in all states seem to be undergoing similar trends, we then tested the following predictions regarding comparisons between California and the other 10 western states.

### Prediction: After 15 years of intensive puma control, states with sport hunting of pumas should experience higher deer densities and deer kill densities of deer by hunters than California

We compared California and the 10 states to determine whether or not either deer densities or kill densities changed from the onset of higher puma kill rates over most states in 2000 to 2015. We found that most states had lower deer and deer kill densities (Fig. 10). Of the states that had positive changes in deer and deer kill densities, California ranked 2^nd^ and 3^rd^ highest respectively (Fig. 12a & b). For most states, deer densities and kill densities have been gradually declining in spite of record high kill rates of pumas. These results do not support the prediction that the intensive killing of pumas through the sport hunt has led to increased deer numbers over the last 15 years.

**Fig. 10.**
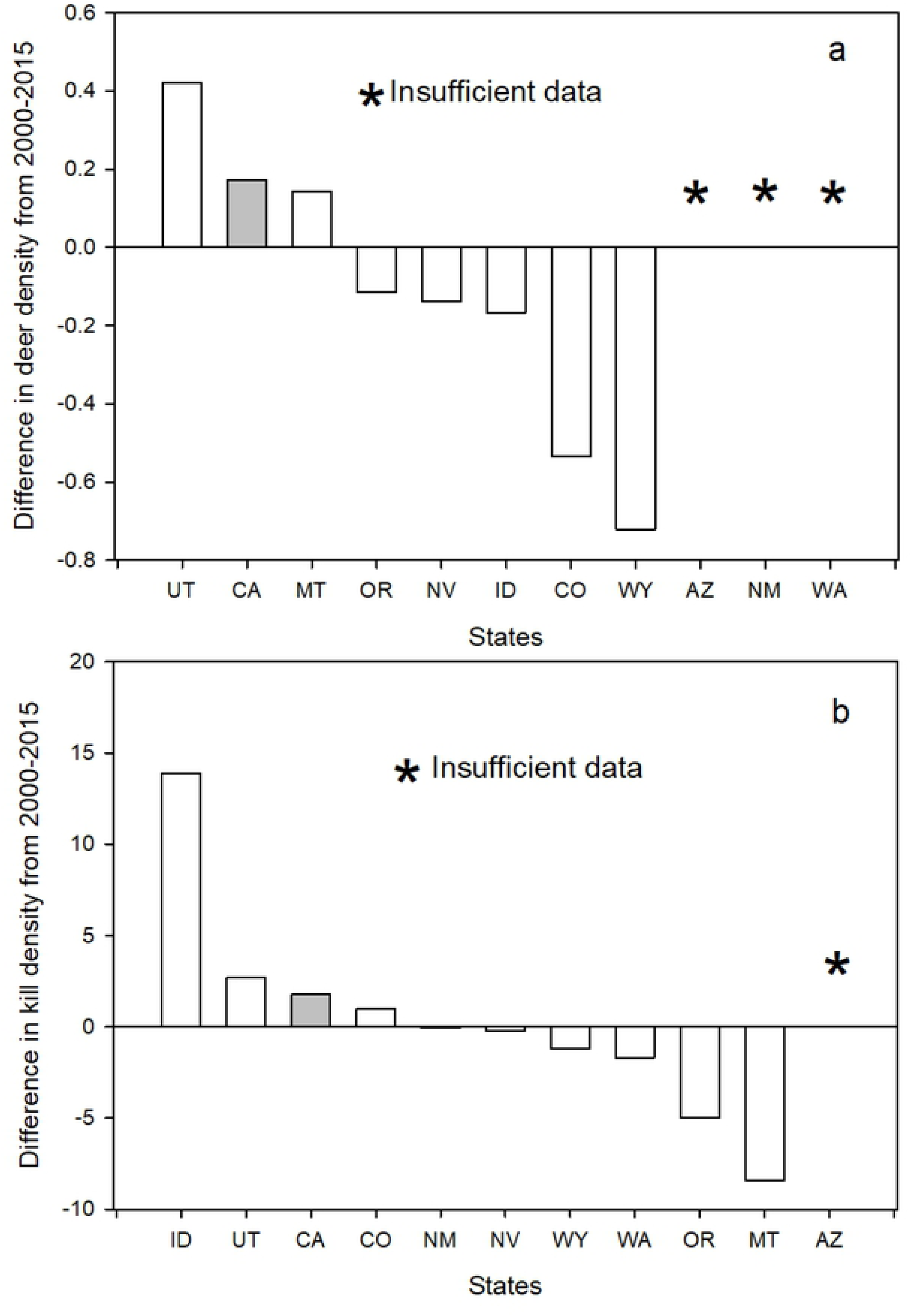
(a) Difference in deer density (#/km^2^) from 2000 to 2015 for California and 7 of the 10 states with a sport hunt on pumas where data were available. Figure 10b difference in kill density (#/100 km^2^) from 2000 to 2015 for California and 9 of the 10 stats with a sport hunt on pumas. There were insufficient data from Arizona for this analysis. States are identified by their standard two letter postal codes.

**Fig. 11.**
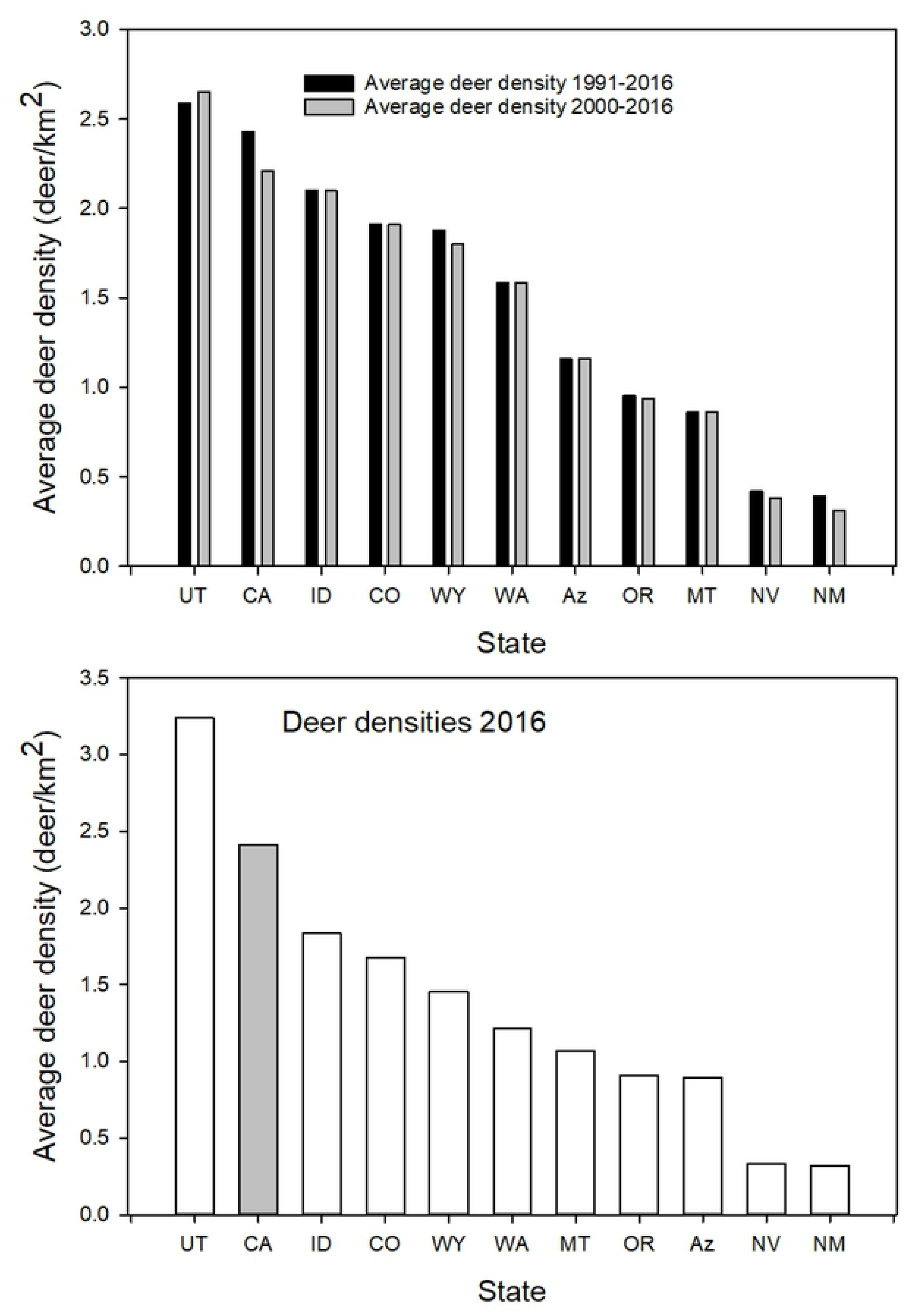
(a) Ranking of mean deer densities (deer/km^2^) from 1991-2016 and 2000-2016 for California and the 10 states with a sport hunt on pumas. Figure 11b is the mean deer densities in 2016 for California and the 10 states with a sport hunt on pumas. States are identified by their standard two letter postal code.

**Fig. 12.**
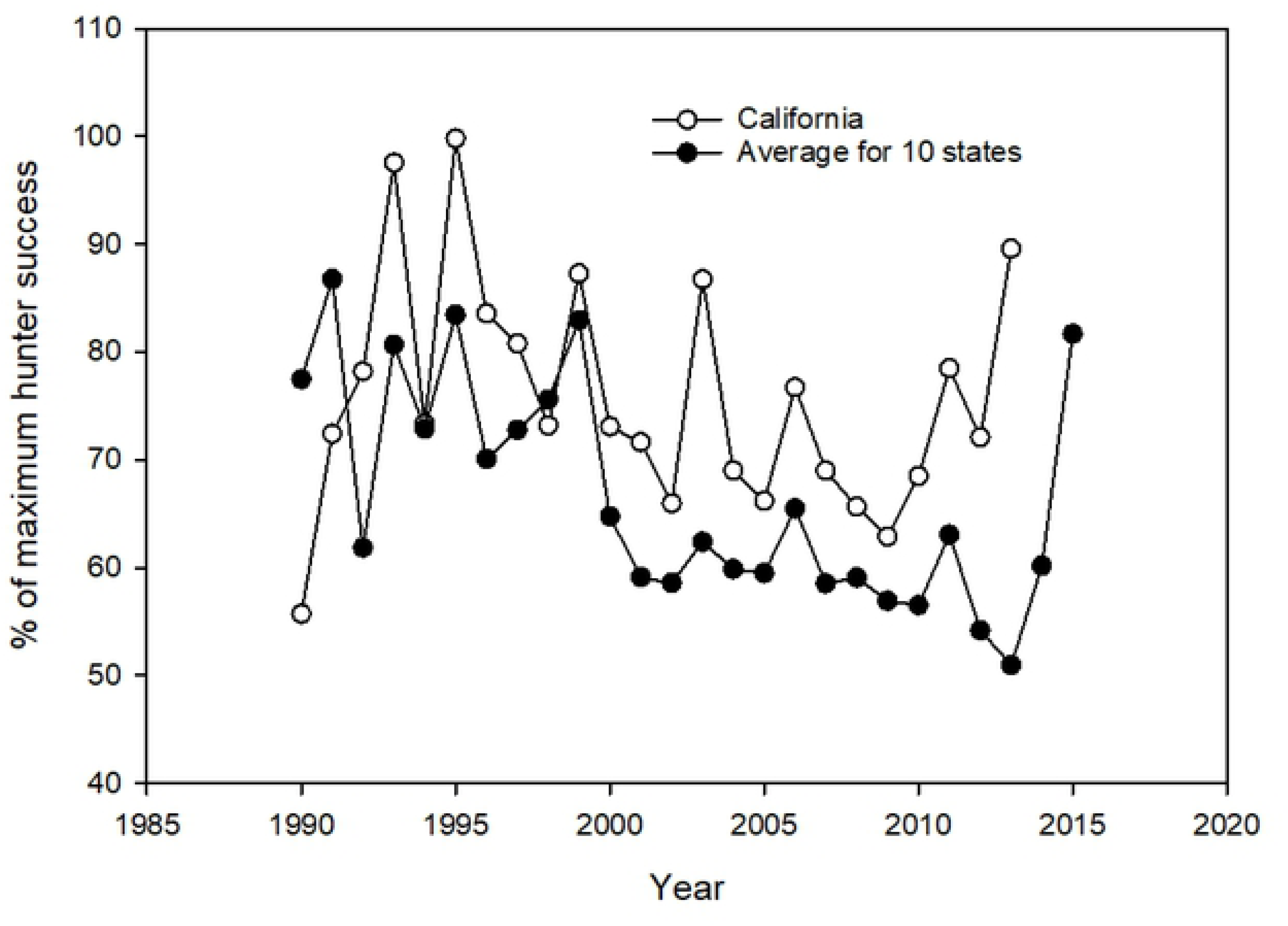
Annual percent hunter success expressed as a percentage of the year of the highest hunter percent success for California and the 10 states with a sport hunt on pumas. The curve for the 10 states is the mean of these states’ values.

The primary prediction regarding deer is that the higher levels of killing of pumas should result in higher deer densities. We compared average deer densities among the 11 states for 1990-2015 and 2000-2015 (Fig 11a) and 2016 (Fig 11b). Among the 11 states, California had the second highest deer densities in all three time periods.

We further correlated both deer density and deer kill density with puma kill densities for the previous year for the 11 states (Table 1). The prediction was that increasing numbers of pumas killed should have a positive effect on deer density and the number of deer that hunters killed. In all cases except one (Washington), deer density and deer kill density either did not significantly correlate with puma kill densities or were negatively related, i.e. higher number of pumas killed resulted in lower deer densities and kill densities (Table 1).

**Table 1:**
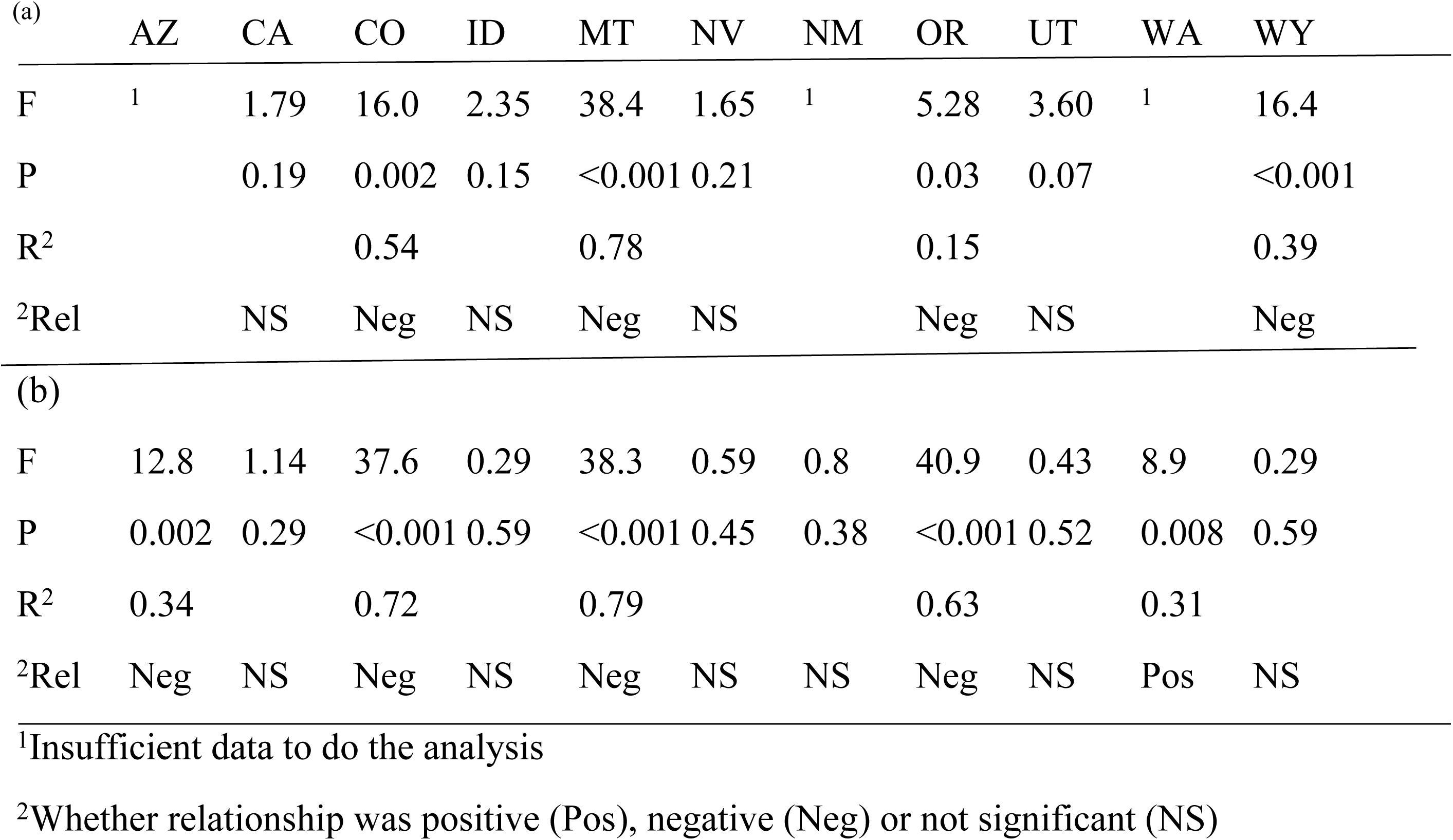
Correlations of deer density estimates and the number of pumas killed the previous year. Data are for the 11 western states (1a) and the deer numbers killed by hunters (deer kill density) and number of pumas killed the previous year (1b). Data are from 1990 to 2015.

|  |  | AZ | CA | CO | ID | MT | NV | NM | OR | UT | WA | WY |
| --- | --- | --- | --- | --- | --- | --- | --- | --- | --- | --- | --- | --- |
| 503 | F | <sup>1</sup> | 1.79 | 16.0 | 2.35 | 38.4 | 1.65 | <sup>1</sup> | 5.28 | 3.60 | <sup>1</sup> | 16.4 |
| 504 | P |  | 0.19 | 0.002 | 0.15 | <0.001 | 0.21 |  | 0.03 | 0.07 |  | <0.001 |
| 505 | R<sup>2</sup> |  |  | 0.54 |  | 0.78 |  |  | 0.15 |  |  | 0.39 |
| 506 | <sup>2</sup>Rel |  | NS | Neg | NS | Neg | NS |  | Neg | NS |  | Neg |
| (b) |  |  |  |  |  |  |  |  |  |  |  |  |
| 508 | F | 12.8 | 1.14 | 37.6 | 0.29 | 38.3 | 0.59 | 0.8 | 40.9 | 0.43 | 8.9 | 0.29 |
| 509 | P | 0.002 | 0.29 | <0.001 | 0.59 | <0.001 | 0.45 | 0.38 | <0.001 | 0.52 | 0.008 | 0.59 |
| 510 | R<sup>2</sup> | 0.34 |  | 0.72 |  | 0.79 |  |  | 0.63 |  | 0.31 |  |
| 511 | <sup>2</sup>Rel | Neg | NS | Neg | NS | Neg | NS | NS | Neg | NS | Pos | NS |

<sup>1</sup>Insufficient data to do the analysis
<sup>2</sup>Whether relationship was positive (Pos), negative (Neg) or not significant (NS)

### Prediction: There should be a positive correlation between deer hunter success and the sport killing of pumas the previous year

Hunter success is a common metric used by game agencies to judge the success of providing deer hunting opportunities to hunters. Hunter success, which can differ widely over large geographic areas such as states, is influenced by various factor. These factors, which include but are not limited to deer density, season length and type (e.g. bucks only or either sex), weather, and how hunter success is calculated (e.g. total deer licenses sold versus “active” hunters in the field (Wyoming data)), make useful across state comparisons unrealistic. A further complication is that game agencies calculate how many deer are killed in different ways, e.g. mandatory check-ins vs surveys. To analyze trends within states, we compared these data separately within the 11 states. As with deer densities, we found no correlation or in the cases of Oregon (F = 15.5, P < 0.001, R^2^ = 0.41) and Wyoming (F = 16.9, P < 0.001, R^2^ = 0.55), negative correlations, i.e. higher kill levels of pumas were associated with lower hunter success. These results indicated that the level of puma mortality did not produce the desired effect of higher hunter success.

To make comparisons between California and the other 10 states regarding the pattern of hunter success over the 1990-2015 timespan, we calculated the percentage each year’s hunter success was to the year the maximum hunter success was recorded (See Methods). We then averaged the percentages for the 10 states and plotted the results with the data from California (Fig. 12). As can be seen in Fig. 12, though the amplitude of the percent maximum for each year was different at times, the patterns of increases and decreases in hunter success appeared quite similar. Most years when hunter success went up in the ten states, it also did in California and vis versa. This indicates an underlying common factor other than puma predation could be driving hunter success.

Another metric we used to ascertain if the killing of pumas by sport hunting was having a positive impact on deer availability for human hunters was the estimate of the number of deer per hunter available in the state. Recall that the prediction was that if sport hunting of pumas was having a positive effect, then we should see 1) a higher average number of deer per hunter in the 10 hunting states compared to California over the 1990-2015 timeframe, 2) the 10 states with a sport hunt should have increases in deer per hunter estimates from 2000-to 2015 (there were insufficient data from several states for the 1990-2015 comparison), and 3) the kill level of pumas within a state should have a positive correlation with the number of deer per hunter.

In the first comparison, 5 hunting states had more and 5 had fewer deer per hunter than California (Fig. 13a). A one sample t-test comparing the 10 sport hunting states with California indicated no statistical difference. In the second comparison, after 15 years of puma mortalities, 6 states, including California reported a decline in deer per hunter, with California having the smallest decrease, whereas two states (Utah and Oregon) reported more deer per hunter (Fig. 13b). In the third comparison, correlating the number of deer per hunter for a given year with the density of puma kills the year before yielded two significant correlations, Oregon had a positive correlation (F = 31.5, P < 0.001, R^2^ = 0.59) and Wyoming had a negative one (F = 8.8, P = 0.014, R^2^ = 0.42). The remaining states, including California had no significant relationship between puma kill levels and the number of deer per hunter within their borders.

**Fig. 13.**
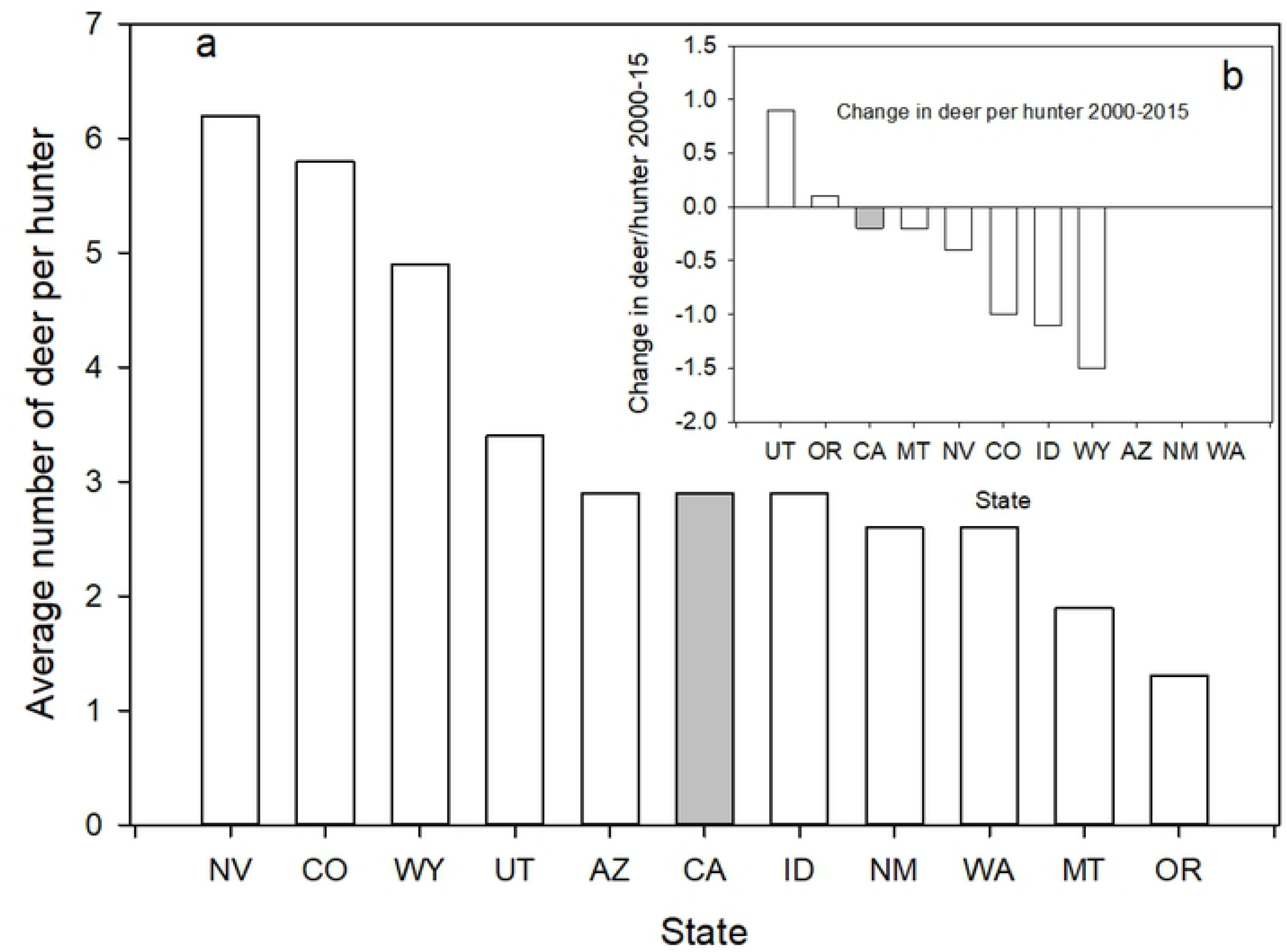
(a) Mean number of deer per hunter (number of deer/number of hunters) for California and the 10 states with a sport hunt on pumas. Figure 13b is the change in the number of deer per hunter from 2000 to 2015 for California and 7 of the 10 states with a sport hunt on pumas. Data were not available for Arizona, New Mexico, and Washington to make this comparison. States are identified by their standard two letter postal codes.

## Discussion

Sport hunting has been widely employed by state wildlife agencies in the western United States to manage puma since the early 1970’s. Stated agency justifications for this practice are based on the hypotheses that widespread killing of puma by hunters will suppress puma numbers, thereby reducing undesirable puma impacts on human safety, livestock, and ungulate populations (e.g. https://idfg.idaho.gov/wildlife/predator-management, Accessed on February 28, 2018). This management strategy has been used by 10 westerns states since the early 1970’s to kill increasing numbers of puma. There has now been sufficient time to test whether sport hunting is having the desired effects relative to an un-hunted puma population, i.e. California. By making various comparisons between the 10 sport hunting states and California we tested the hypotheses that a sport hunt would: 1) suppress puma numbers at levels lower than would be expected without a sport hunt and subsequently, 2) reduce problematic puma-human interactions, 3) reduce puma depredation on domestic livestock, and 4) reduce the impact of puma predation in wild ungulate numbers.

Within the constraints of the robustness of the data available, we found no evidence those data support the hypothesis that sport hunting has long-term effects on puma numbers. California reports similar average densities of pumas as the 10 hunting states after 40+ years of increasing sport hunting rates by those states (Fig. 2). These results concur with those of [14] who found no evidence of sport hunting having a regulating impact on puma populations. In their study, a main factor possibly negating any controlling influence of hunting was the immigration of dispersing individuals from surrounding areas [35]. As any dispersing pumas from California would only have a limited regional impact, e.g. Arizona, Nevada and Oregon, it is unlikely that dispersing individuals from California are affecting puma abundance across the entire West.

Additionally, in Oregon the records indicate that increasing killing of pumas is associated with increases, not decreases, in estimated puma numbers (Fig 3). As reported by state the agency, both puma numbers and puma kill rates in Oregon have substantially risen over the last 20 years, contrary to what would be predicted by the sport hunting model. Consequently, based on their own data, this alone would argue against further use of sport hunting of pumas as a management tool.

Pumas are widely recognized by wildlife managers as one of the more difficult species to enumerate. Most state agencies admit their population estimates for pumas have low reliability; Idaho, does not attempt to estimate puma numbers. Nevertheless, such estimates are often used to justify management decisions to increase the number of puma killed through sport hunting. However, we found no evidence, within the sensitivity of the data collected to distinguish differences, that the sport hunt has had the desired effect of reducing puma abundance.

The data from California appears to support some of the original studies proposing that social organization of pumas is a limiting factor on total puma abundance [36]. However, in the absence of sport hunting, the number of pumas in California is probably regulated by a combination of social organization and prey abundance [25, 26]. Puma populations fluctuate with prey abundance [25] and when prey abundance is low, its availability probably limits the number of pumas an area can support regardless of social limitations. However, with higher prey levels and increasing puma numbers, social strife possibly sets the upper limit of puma densities in an area, apparently regardless of whether the population is hunted or not. Recent work on social organization in pumas [37] indicates even more complex social interactions than earlier thought. These interactions underscore the importance of a stable social structure that sport hunting appears to disrupt [38].

Regarding the prediction that sport hunting of puma should reduce risk to human safety, recent studies have indicated that the use of this management tool may have just the opposite effect [20, 39]. The results of our multi-state analysis in general supports the more regional findings in that first, there appears to be no relationship between sport hunting of pumas and human safety/conflicts. For each timeframe since 1972 considered, California has similar total numbers of recorded puma attacks as some hunting states and the third lowest number of per capita attacks (Fig. 4). In our calculation of per capita rates, we considered the total populations of each state. This was in part because of the difficulty in separating out urban and rural population numbers but also in recognition that in many of the states, pumas are widespread throughout the states and readily use suburban and exurban areas [40, 41]. This is especially the case for California where pumas are commonly reported near and in major housing developments [41, 42, 43, 44]. Though Florida was not included in this analysis, it should be noted that Florida panthers are totally protected, living in one of the most densely human populated area of the U.S. and there have been no attacks on humans over the same time intervals considered [45].

Contrary to predictions, higher kill rates of puma coincided with higher numbers of incidents in two of the three states where data were available, Utah and Washington. Our results from Washington from 1992-2015, concur with a 5-year analysis (2005-2010) of that state [20] and a more recent analysis from British Columbia [39]. Indiscriminate killing of pumas appears to disrupt social structure and stability [37, 38], resulting in younger less experienced individuals having more conflicts with humans [20].

The risk of puma attacks on humans is normally extremely low (approximately 2/year across the 15 states where pumas are found). This is in comparisons to normally excepted higher risks from other wildlife species, e.g. 150-200 human fatalities per year in deer-car collisions [46]. As sport hunting of deer is not used to address these higher incidences, we found no justification for the rationale to use sport hunting pumas to address human safety concerns.

The western states we considered all have major extensively managed livestock operations where livestock, mainly cattle and sheep, are grazed on open pasture, often in the same habitats used by pumas. Pumas do prey on these livestock. However, as with human risks, the average rate of depredation is low, especially when considered as a percentage of the total number of head of livestock exposed to the risk of puma predation. This being the case, however, it is still valid to ask: could the sport hunt of pumas further lower the predation rate on cattle and sheep? Based on our analysis of the data, the answer appears to be no. Comparing the 10 puma hunting states to California we found no difference in the percent loss of total inventory of cattle (Fig. 5) or sheep (Fig. 7). We also found no effect of puma kill rates among all the states and percentage of inventory lost (Fig 6). On the contrary, in concurrence with data from Washington (20) (Peebles et al. 2013), we did find higher percentages of calves killed by pumas with higher puma kill rates in Oregon (Fig. 6b) and a similar response for sheep and lambs in Utah. Peebles et al. [20] credited the higher rates of livestock predation in their study to the disruption of the social order by the indiscriminate killing of resident individuals by the sport hunt. It would appear that in these two states at least, a similar social upheaval might be occurring. In conclusion, again, our multi-state analysis failed to demonstrate any reduction of livestock depredation attributable to the sport hunt of pumas.

The last prediction we tested was whether the sport hunt of pumas resulted in “more game in the bag” for deer hunters. Much to the frustration of game agencies, rising and falling deer populations seems to be the norm for most of the western states [18, 47]. Over the long term, the general pattern based on available data has been a significant increase in deer numbers after deer were protected from uncontrolled hunting prior to the 1920’s (Fig. 8). It appears that in most states, including California, deer populations peaked around 1960 and then declined dramatically after, with a minor recovery in the mid 1980’s. There have been innumerable number of studies and several reviews of those studies to try and identify what is driving deer populations. The usual suspects have been considered extensively, e.g. weather, habitat destruction, over-browsing, and predation. Many studies have tested whether pumas are affecting deer populations [17, 26, 48, 49] and at least three reviews of these studies exist [15, 16,18]. The general consensus is that pumas are not affecting deer numbers and killing puma only will enhance deer populations under very limited circumstances in space and time [15,16, 49]. Similar non-impacts by pumas have been found for elk [48, 50]. Yet, most agencies still use blanket killing of pumas by sport hunting over most of their state, an approach which appears unjustified. In one study [25, 49] puma population numbers were monitored through the increase in deer numbers in the mid 1980’s and their subsequent decline. Based on the demographics of the puma population [25], it appeared that deer numbers were more likely driving puma numbers, with deer numbers being more affected by weather conditions [49]. Our multi-state analysis in general concurs with these many studies and reviews.

We first found that average annual deer densities in California were the second highest for all time intervals considered (Fig. 11). The differences in deer densities among states could be due to inherent limits in habitat carrying capacity. This is possibly the case for the states of Arizona, Nevada, and New Mexico as they encompass primarily desert environments. However, most other states did have some years that equaled or exceeded the average deer densities for California. This indicated that while they had the potential to have similar or higher densities, the sport killing of puma did not seem to lead to those higher densities. We found only one state, Washington, where deer densities were positively related to the number of puma killed (Table 1). In the other states, including California, there was either no relationship or it was a negative one, e.g. lower deer density with higher number of puma killed. Additionally, deer densities in most states, including Washington, have decreased in association with the higher levels of puma kill rates since the year 2000 (Fig. 10).

A major management goal of most agencies is to provide hunters with a reasonable level of success. That success can be measured, in part, by the number of deer per hunter. Regardless of the total deer density, the more deer per hunter, the more likely a hunter can be successful. This can be seen in New Mexico where, though it had the lowest deer density of all the states (0.4 deer/km^2^), had the highest deer per hunter (6.2) and thus had a relatively high hunter success rate (42.2%). California had an equal to or higher number of deer per hunter as 7 of the sport hunting states, indicating that there were similar numbers of deer available to hunters in most states regardless of whether pumas were removed. Further, when we compared the change in deer per hunter data for each state after 15 years of intense killing of puma, most states had fewer deer per hunter, with California having the least decline.

The overall conclusion of these comparisons is a rejection of the sport hunting hypothesis regarding 1) suppression of puma numbers, 2) reduction of problematic puma-human encounters, reduction of puma predation on livestock depredation, and 4) reduction of the impact of puma predation on wild ungulate populations. The results of these comparisons concur with a growing number of regional studies that find no consistent evidence that sport hunting is functioning as an effective management tool. It may, in fact, be having the opposite results [14, 20, 39]. It is becoming evident that under the guidelines of adaptive management, in the absence of evidence of its efficacy, state agencies should refrain from prescribing sport hunting as a management tool.

Whether or not sport hunting of pumas should be continued as a hunting opportunity to hunters is, however, a decision that should be made through the democratic process and involve all the citizens within each state. As specified by the North American Model of Wildlife Conservation, hunting laws should be created through the public process and should follow the tenets of the NAM that state 1) wildlife is held in the public trust, 2) wildlife use is allocated by law, 3) wildlife should be killed only for a legitimate purpose, and 4) science should be the basis of all decisions [8] (Organ et al. 2012). In making that decision, game agencies will have to justify to the public that maintaining a sport hunt on pumas to solely provide trophy hunting opportunities to a small percent (< 0.2%) of the public is a legitimate reason for killing pumas. They should not use the four proposed outcomes analyzed here as a justification for the continuation of sport hunting of puma. Their own management data just does not support it.

## Acknowledgements

We thank Harley Shaw for providing insightful suggestions on this manuscript. We also acknowledge the immense effort of the hundreds of state and federal employees and members of conservation NGOs who collected and compiled the vast amount of data used in our analyses. It is through their efforts that such a large-scale analysis was possible.

## Supporting information

**S1 Fig. Correlation of number of puma-human incidences reported with puma removal density (#/10,000 km^2^).** Data are for California and the three states (Oregon, Washington, and Utah) for which these data were available.

